# The grass that built the Central Highland of Madagascar: environmental niches and morphological diversity of *Loudetia simplex*

**DOI:** 10.1101/2023.09.25.559324

**Authors:** Tchana O. M. Almary, Joseph White, Fitiavana Rasaminirina, Jacqueline Razanatsoa, Caroline Lehmann, Mijoro Rakotoarinivo, Hélène Ralimanana, Maria S. Vorontsova, George P. Tiley

**Affiliations:** Kew Madagascar Conservation Centre, Antananarivo, MDG; Department of Plant Biology and Ecology, University of Antananarivo, Antananarivo, MDG; Royal Botanic Gardens Kew, Richmond, TW9 3AE, UK; Département Botanique, Parc de Tsimbazaza, B.P. 4096, Antananarivo 101, MDG; Tropical Diversity, Royal Botanic Garden Edinburgh, Edinburgh, EH3 5LR, UK; School of GeoSciences, University of Edinburgh, Edinburgh, EH9 3FF, UK

**Keywords:** Poaceae, grasslands, savanna, species distribution modelling, environmental niche, tropical conservation

## Abstract

1) Research Aims — *Loudetia simplex* is a common and dominant species throughout grassland ecosystems in mainland Africa and Madagascar. It is highly polymorphic, often classified as two taxa endemic to Madagascar: *L. simplex* subsp. *stipoides* and *L. madagascariensis*. A better understanding of the inter- and intra-specific variation between these taxa and its contributing environmental factors could improve our understanding of the history of Madagascar’s grasslands.
2) Methods — The taxonomic status of *L. simplex* subsp. *stipoides* and *L. madagascariensis* was evaluated by morphometric analyses of 119 herbarium specimens. Species distribution modelling was used to determine the most important environmental factors underlying *the L. simplex* distribution in Madagascar versus other African grasslands. We investigated if *L. simplex* in Madagascar could be predicted by distributions across mainland Africa with niche overlap analyses.
3) Key Result — African and Malagasy species exhibited variation potentially associated with environment. Specimens from northern and western Madagascar were taller with smaller spikelets than those from Southern Africa and central Madagascar. *Loudetia simplex* typically occurred in cooler temperatures with high precipitation and pronounced seasonality, but taller populations were found in warmer conditions. Projecting ecological niches of Southern Africa and East Tropical Africa onto Madagascar demonstrates much of the present distribution in the Central Highlands is expected from other natural African grasslands.
4) Key Point — Malagasy and African individuals represent a single species, and the Malagasy species can be considered as a synonym of L. simplex. Distribution models are congruent with pre-human presence of grasslands in Madagascar.

**SOCIETAL IMPACT STATEMENT:** Understanding dominant species like *Loudetia simplex* is necessary to understand the fire-driven grassy ecosystems they create. In Madagascar, grasslands are considered a low-value ecosystem despite their unique biodiversity and crucial importance as zebu rangeland. Only 1.8% of Madagascar’s grasslands are protected despite facing similar threats to forests and biodiversity loss. This study of common grasses will support the management of protected areas by providing information on resource management of vulnerable open ecosystems in Madagascar. Distribution models of common grass species and clear taxonomic classification can help land management stakeholders identify natural grasslands versus degraded forests.

## INTRODUCTION

Madagascar is a biodiversity hotspot (Myers et al., 2000), characterized by its unparalleled floral and faunal endemism (Goodman, 2018, Ralimanana 2022). There are more than 11,000 described plant species on the island (Antonelli 2022), of which approximately 82% are endemic (Goodman, 2018; Antonelli, 2022). This high rate of endemism is due in part to successive dispersal events from lineages with African or Indo-Pacific origins (Callmander et al., 2011; Crottini et al., 2012; Buerki,et al., 2013; Antonelli et al., 2022), despite Madagascar’s long isolation from present-day mainland Africa during the Early Cretaceous (ca. 120 Ma ago) and present-day India during the Late Cretaceous (ca. 85 Ma ago; Ali and Aitchison 2008). Madagascar is presently separated from southeast Africa by the Mozambique Channel, and thus the two landmasses share some macroecological features in addition to species. While both regions have important humid forests, they are dominated by open, grassy ecosystems. The grasses of Madagascar’s Central Highlands and their comparison to mainland Africa have received little taxonomic attention. An improved understanding of the taxonomy and distributions of some common species could help place the natural history of Madagascar’s Central Highlands in the context of other African open grassy ecosystems and facilitate biodiversity investigations, as there is uncertainty in the extent of pre-human grasslands in Madagascar being the centre of a conservation debate (e.g. Joseph et al., 2022a),

Madagascar’s grasslands comprise 65% of the island area (Vorontsova,2016; Lehmann, 2021) and they demonstrate levels of species richness and endemism comparable to the well-recognized ancient grasslands of South Africa (Bond et al., 2008). *Loudetia simplex* (Nees) C.E.Hubb. (Poaceae: Panicoideae: Tristachyideae) is a common and dominant species of the grasslands of Africa and Madagascar, where it is widely distributed in both localities. The species likely dispersed from mainland Africa to Madagascar between 3 and 8 Ma ago (Hackel et al., 2018), consistent with global C_4_ grassland expansions (Edwards and Smith 2010). Previous population genetic analyses with microsatellites suggests there was little to no post-isolation gene flow between Madagascar and South Africa (Hagl et al., 2020), showing recent anthropogenic dispersals do not explain *L. simplex* distributions in Madagascar. Thus, an examination of *L. simplex* taxonomy and distributions would provide insightful with respect to the evolution of Madagascar’s Central Highland grasslands.

The taxonomy of this species is unclear in many cases. In Madagascar it is known under two different names: *Loudetia simplex* subsp. *stipoides* (Hack.) Bosser and *Loudetia madagascariensis* (Baker) Bosser, which is endemic to Madagascar (Bosser, 1966; 1969). Since the assembly of the broader concept of *L. simplex* by Clayton (1974), there have been 29 synonyms of *L. simplex* across the African continent (POWO, 2023). Morphologically, *L. simplex* is a perennial caespitose bunchgrass, up to 150 cm of height, with woolly sheath-leaves base forming a tomentose structure. It is characterized by a fringe of hairs ligule, flat or convolute leaf blades, inflorescence in panicle, spikelet formed by lower and upper glume with two florets, with a bidentate callus subtending the upper fertile floret (Bosser 1969; Fish et al., 2015; Vorontsova et al.,2018; Fig. 1a). This species grows in different habitats and has a phenotypic expression that likely varies with environment (Fish et al., 2015; Bosser 1969). It grows across a range of elevation between 100 m and 2000 m on soil that is sandy, rocky, ferrallitic, eroded, or covered by a crust with poor drainage (Bosser 1969, Solofondranohatra, 2020; Fig. 1b). *Loudetia simplex* is resistant to environmental disturbance and is considered to be a fire-adapted species (Solofondranohatra, 2020).

**Figure 1.**
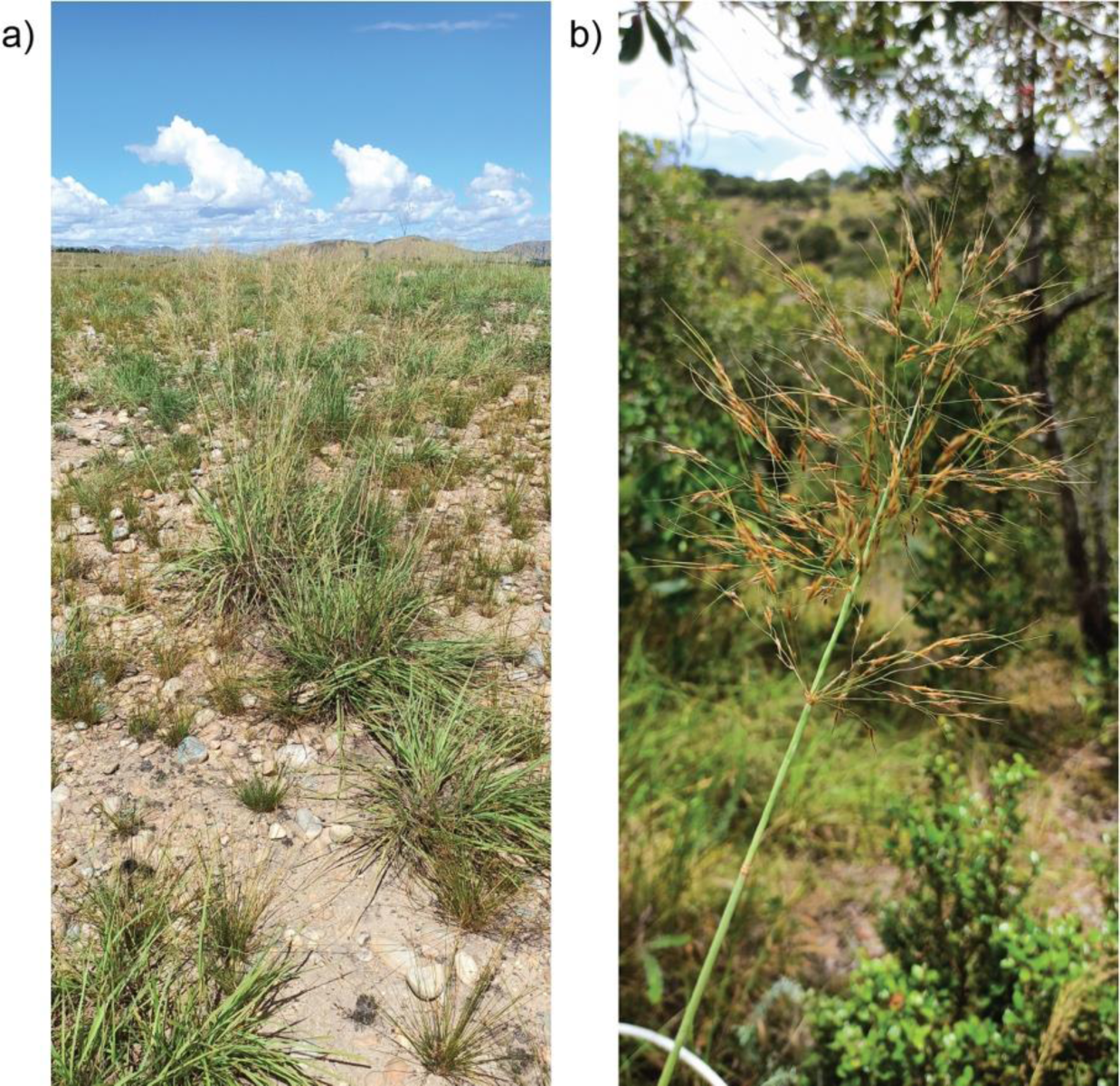
*Loudetia simplex* from Madagascar. a) Habit and habitat of *L. simplex* from western Madagascar (Photo Credit: Tchana O. M. Almary). b) Single inflorescence characteristic of *L. simplex* (Photo Credit: Tchana O. M. Almary).

Given the ecological importance of *L. simplex* across African grasslands and its taxonomic uncertainty within Madagascar, we revisit the two subspecies hypothesis (Bosser 1969) and aim to develop a better understanding of Malagasy grasslands by comparisons of *L. simplex* across Africa. In this study, we used analyses of morphological and environmental variation from Malagasy and mainland African *L. simplex* specimens to address the following questions: 1) Are Malagasy and mainland African *L. simplex* the same species?, 2) Are there subspecies within Madagascar?, 3) What are the environmental variables underlying *L. simplex* distributions in Madagascar and mainland Africa?, and 4) Can Malagasy *L. simplex* dominated grasslands be predicted by distributions from other African grasslands?

## MATERIALS AND METHODS

### Morphological Approach

#### Sampling and study regions

119 *Loudetia simplex* specimens were used to compare African and Madagascan individuals (Supplementary Table S1). 36 specimens were measured at the Parc Botanique et Zoologique Tsimbazaza herbarium (TAN) and 63 primarily mainland African specimens at the Royal Botanic Gardens, Kew herbarium (K). Measurements were also made on 10 new specimens collected in western Madagascar. Three herbarium specimens were excluded from downstream analyses due to a lack of leaves or incomplete samples with no leaf sheath. Specimens were chosen to reflect the broadest possible distribution of open grassy ecosystems where *L. simplex* is dominant (Fish et al., 2015; Hagl,2020). Specimens were sorted into the following populations *a priori* for hypothesis testing: Southern Africa (SA), South Tropical Africa (STA), East Tropical Africa, Western Madagascar (WM), Southern Madagascar (SM), Central Madagascar (CM) and Northern Madagascar (NM) (Fig. 2). Groupings of mainland African populations are based on the World Geographic Scheme for Recording Plant Distributions (Brummitt, 2001).

**Figure 2.**
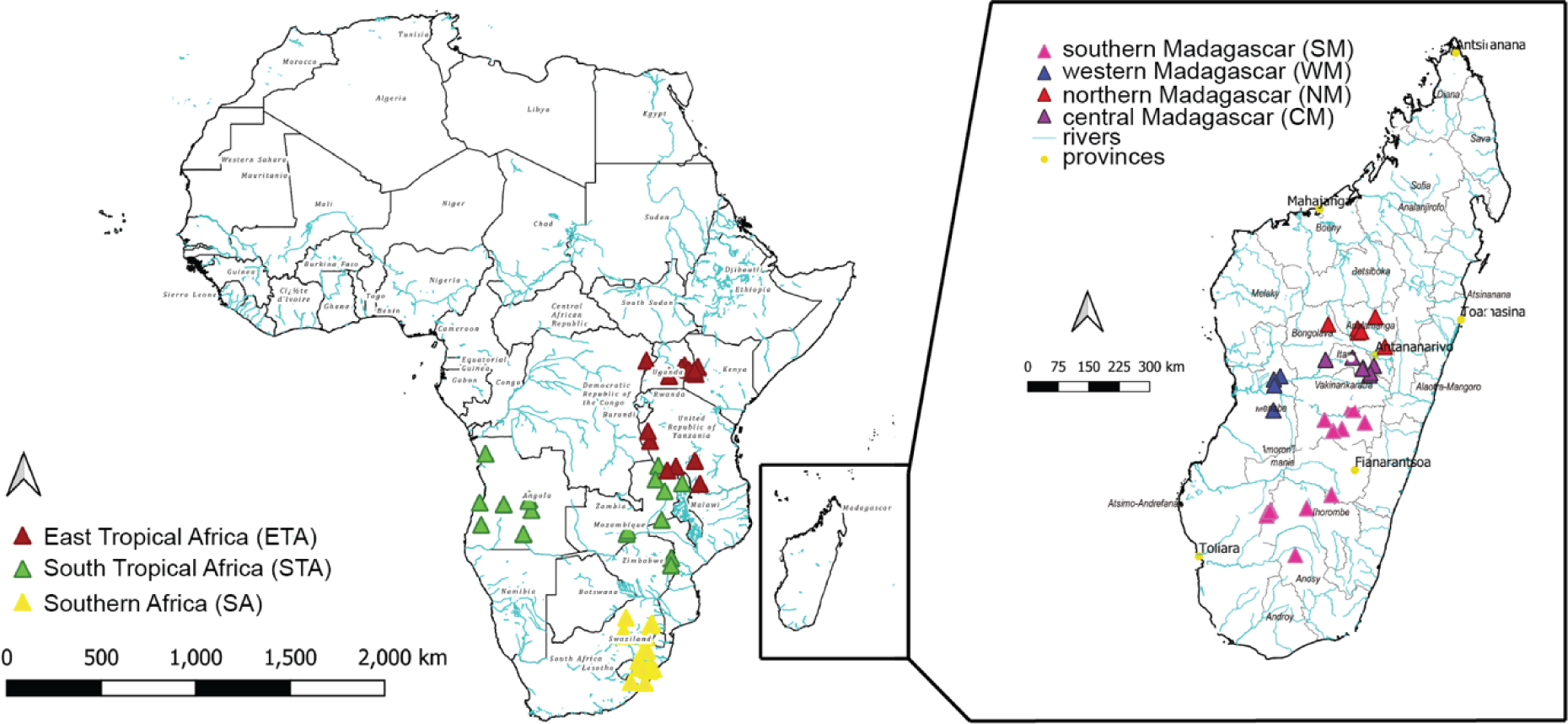
Distribution of measured herbarium specimens. Distribution of individuals used for morphological analyses and their regional assignments.

#### Morphological data

We measured 16 traits characterized by Bosser (1969), which included scoring 10 qualitative characters (culms profile, leaf sheath character, leaves form, inflorescence colour, inflorescence density, hairiness, spikelet size and form, lower glume tips and callus form). Our six quantitative variables (height, leaf length, inflorescence length, spikelet length, lower glume length and awn length) were measured under a dissecting scope as necessary. Because we are studying potentially a single complex species, many qualitative characters were found to be constant (Supplementary Table S1), and only four of the quantitative characters were selected for statistical analyses. Character descriptions are covered in Table 1.

**Table 1.** Traits use for morphological analyses.

| **Parameter** | **Abbreviatio**<br>**n** | **Variable**<br>**Type** | **Encoding or Measurement Unit** |
| --- | --- | --- | --- |
| Height | H | QT | cm |
| Culms | CM | QL | slender: 0 , large: 1 |
| Base of leaf sheath | LS | QL | tomenteous: 0 , fibrous: 1 |
| Leaf length | LL | QT | cm, in an interval |
| Leaf form | LF | QL | large: 1, filiform: 0 |
| Color of<br>inflorescence | COI | QL | brown: 0, beige: 1, darkbrown: 2,<br>gold: 3 |
| Character of<br>inflorescence | CHI | QL | open: 0, contracted: 1, big: 0,<br>small: 1 |
| Inflorescence length | IL | QT | cm |
| Size of spikelet | SS | QL | big: 0, small: 1 |
| Profile of spikelet | PS | QL | thin: 0, large: 1 |
| Hairiness | HA | QL | Glabrous: 0, Hairy: 1 |
| Spikelet length | SL | QT | mm |
| Lower glume length | LGL | QT | mm |
| lower glume tip | LGT | QL | acute: 0, rounded: 1 |
| Awn length | AL | QT | mm |
| Callus | CA | QL | bidentate: 0, oblong: 1 |

†QL = qualitative, QT = quantitative

#### Statistical analyses

All statistical analyses and visualization were conducted in R studio using R v.4.2.0 (R core team, 2022). We used one-way analysis of variance (ANOVA) analyses to test for differences in morphological characters between Malagasy and mainland African collections as well as among populations. A post-hoc Tukey’s honestly significant difference (HSD) test was used to identify significantly difference pairs assuming an α of 0.05. A principal component analysis (PCA) was used with the MorphoTools2 R package (Šlenker et al., 2022) to visualize multivariate differences between Madagascar and mainland Africa.

### Environmental Approach

#### Ecological data

We used a suite of environmental variables to understand and predict the distributions of *L. simplex*. These included: temperature and precipitation data from WorldClim V1 (Hijmans et al., 2005), mean values of climate and climatic water balance data from TerracClimate (Abatzoglou et al., 2018), elevation data from the Shuttle Radar Topography Mission (Jarvis et al., 2008), geomorphology data, including compound topographic index, topographic roughness index and slope from Geomorpho90m (Amatulli et al., 2020) and edaphic data at 0-20 cm from iSDAsoil (Miller et al. 2021), including soil bulk density, effective cation exchange capacity, soil organic carbon, total carbon, total nitrogen, clay content, sand content, silt content and pH. TerraClimate data was upsampled from +-5 km to +-1 km, while the SRTM, Geomorph90m and iSDAsoil data were all downsampled to +-1km to match the spatial resolution of the WorldClim dataset. The environmental data were masked to include sub-Saharan Africa below 18.5°N and Madagascar, covering all digitised *L. simplex* occurrences, and were processed using Google Earth Engine (Gorelick et al., 2017). We tested for multicollinearity among the environmental variables across sub-Saharan Africa and Madagascar using Pearson’s R at a 0.7 threshold (Leroy et al., 2015), leaving 11 uncorrelated variables (Supplementary Table S2).

Presence occurrence records were downloaded from Global Biodiversity Information Facility (GBIF, https://www.gbif.org/) and Botanical Information and Ecology Network (BIEN) v4.1 (http://bien.nceas.ucsb.edu/bien/). We added 11 new presence data points from our field trip from western Madagascar in 2022 (Supplementary Table S1). Citizen science data have known issues with inconsistent metadata standards and geographical biases, imprecision, or errors (Zizka et al., 2019). We therefore cleaned the pooled occurrence records by removing duplicates, masking to terrestrial landmasses, and removing occurrences intersecting within a 2 km radius of zoo, herbaria and country capital centroids with the CoordinateCleaner R package (Zizka et al., 2019). Lastly, to further correct for sampling biases in our presence records we applied spatial thinning to include one presence per 1 km^2^ grid cell to match the spatial resolution of our environmental variables with the FLEXSDM v1.3.2 R package (Velazco et al., 2022). For distribution modelling, we produced an equal sample of pseudo-absences to cleaned presences and randomly distributed them outside of a 1 km buffer of each presence point for each model (Velazco et al., 2022).

#### Species Distribution Models

To determine the relative influence of environmental variables on *L. simplex* distributions and predict their habitat suitability, we ran Random Forest (RF) models (Liaw & Wiener, 2002) using the SDMtune R package (Vignali et al., 2020) on four regions, which at least partially matched the sites from our morphological analyses. This included: 1) the full *L. simplex* mainland African and Malagasy range, 2) East Tropical Africa (ETA), which was restricted to Mozambique, Zimbabwe, Zambia, Malawi, United Republic of Tanzania, south-eastern Democratic Republic of the Congo, Burundi, Rwanda, Uganda, and Kenya, 3) Southern Africa (SA), comprised of South Africa, Lesotho, and Eswatini, and 4) Madagascar. Note that we included countries from the South Tropical Africa and West-Central Tropical Africa botanical regions (Brummitt, 2001) in our ETA model, but only use ETA henceforth for brevity. By modelling species distributions based on occurrence and spatial environmental data (Franklin, 2010; Peterson et al., 2011) as a proxy for habitat suitability (Warren & Seifert, 2011), we explored environmental differences between *L. simplex* in Madagascar versus mainland Africa at different scales. Scripts for analyses are available at TOMA’s GitHub (https://github.com/AlmaryT/Loudetia_SDM).

#### SDM training, evaluation, and prediction

To further account for spatial sampling biases of occurrence records, we performed 5-fold spatial cross-validation using environmental blocking to separate training and testing folds of presences and pseudo-absences from the blockCV R package (Valavi, Elith, Lahoz-Monfort, & Guillera-Arroita, 2019). This separates training and testing folds in environmental space across our 11 selected variables to reduce spatial autocorrelation of predictions. This was followed by hyperparameter tuning to find the best-performing models.). We tuned *mtry* (2 to 10), *ntree* (250 to 1500) and *nodesize* (1 to 10) parameters using a random search across 50 possible hyperparameter combinations. The performance of trained RF models was determined using optimal Area Under Curve (AUC; Fielding & Bell, 1997) scores. Final model evaluations are presented using AUC and True Skill Statistic scores. We assessed environmental variable outcomes using a permutation importance approach (across 10 permutations) and marginal response curves where all other variables are set at average values.

Lastly, we explored the niche overlap between *L. simplex* modelled in SA and ETA, the two nearest regions, and projected onto the environmental space in Madagascar. Habitat suitability maps were compared using the respective binary range maps. Thresholds to produce the binary maps were calculated as the mean maximum training sensitivity plus specificity scores across each model fold. All rasters were then aggregated to remove rogue cells using a thresholding approach, removing clumps of cells with less than 1000 pixels. Niche overlap was tested on the 1) full projections from SA and ETA onto Madagascar and 2) projections masked to exclude eastern humid forest regions where *L. simplex* would be shaded out. Madagascar eastern humid forest regions were extract from Moat & Smith (2007) to include the classes labelled ‘humid forest’ and ‘degraded humid forest’. Niche overlap was calculated using Schoener’s statistic (D; Schoener 1968) and a standardised Hellinger distance (I; Warren et al., 2008), which might be more appropriate for comparing distribution models. Both statistics are bounded between 0 and 1, such that 0 is no overlap between niches and 1 is perfect overlap. Overlap statistics were calculated with the dismo v1.3.9 R package (Hijmans et al., 2022).

## RESULTS

### Morphological Variation within Loudetia simplex

First, we analysed the variation of the four quantitative traits between mainland Africa and Madagascar. ANOVA tests suggested significant morphological differences across all traits except height (Fig. 3a; Supplementary Table S3). However, a PCA showed no discernible clustering of individuals (Fig. 3b).

**Figure 3.**
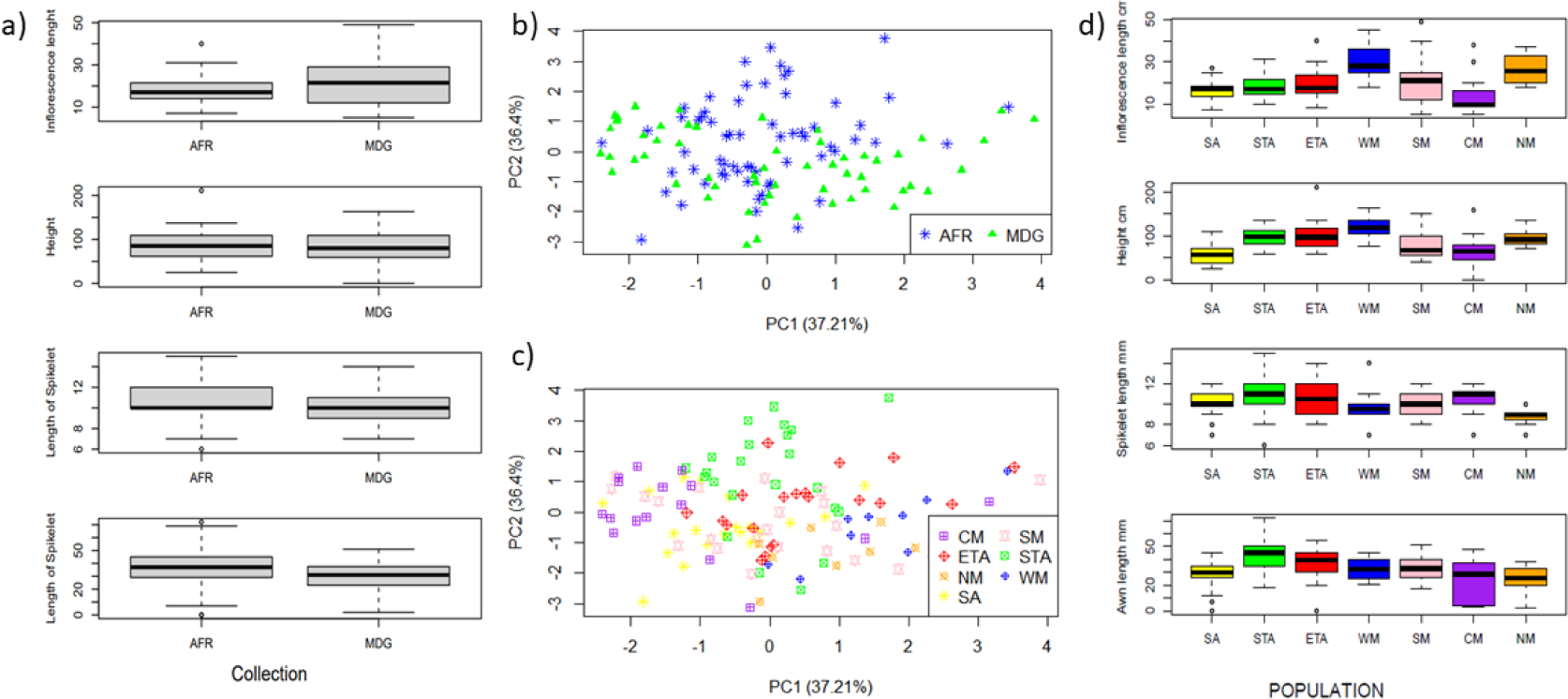
Morphological differences among populations. a) Boxplots of quantitative characters compared between Madagascar and mainland Africa. Significant differences among distributions are given in Supplementary Tables S3 and S4. b) PCA of morphological data for individuals from Madagascar (MDG) or mainland Africa (AFR). c) The PCA with individual data points assigned to regions. d) Boxplots of quantitative characters grouped by *a priori* regions within Madagascar and mainland Africa.

Therefore, we tested for regional differences within Madagascar and mainland Africa. We found significant differences in quantitative characters across regions (Fig. 4a; Supplementary Table S3). For total plant height within mainland Africa, ETA and STA populations were taller than SA (Table 2). Within Madagascar, we found the WM and NM populations were taller than CM and SM (Table 2). However, there was no significant height difference between the ETA and STA populations or the WM and NM populations (Fig. 3c; Supplementary Table S4).

**Figure 4.**
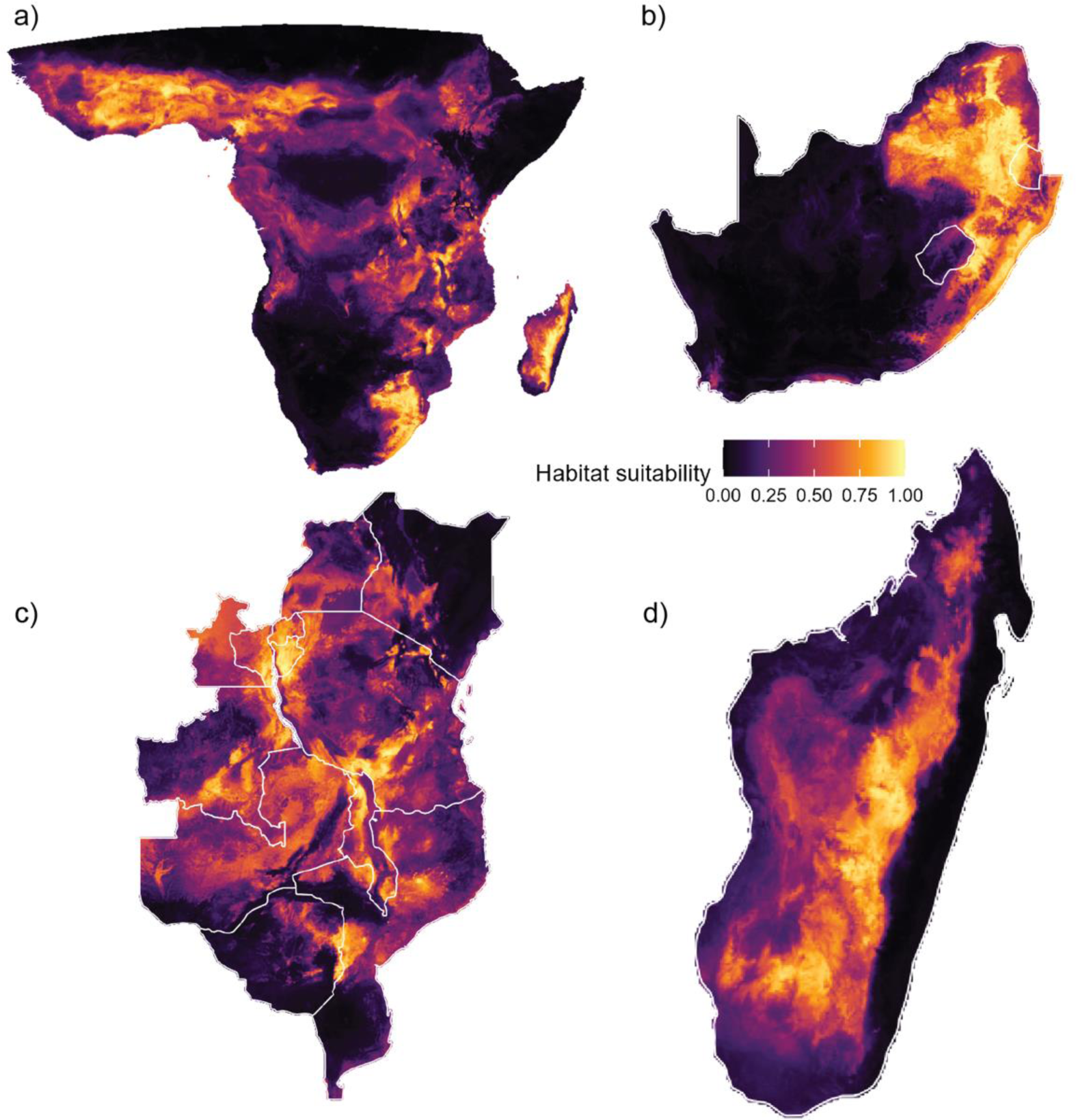
Habitat suitability of *Loudetia simplex*. Habitat suitability determined by independently trained models representing a) Africa, b) Southern Africa (SA), c) East Tropical Africa (ETA), and d) Madagascar.

**Table 2.** Population-level trait summary statistics.

| Population | Sample Size | Plant Height<br>(H) |  | Inflorescence<br>Length (IL) |  | Spikelet<br>Length (SL) |  | Awn Length<br>(AL) |  |
| --- | --- | --- | --- | --- | --- | --- | --- | --- | --- |
|  |  | mean<br>n | variance<br>e | mean | variance | mean<br>n | variance<br>e | mean<br>n | variance<br>e |
| SA | 20 | 58.5 | 545.5 | 16.6 | 27 | 10 | 1.3 | 28.9 | 134.1 |
| STA | 23 | 96.6 | 478.5 | 18.1 | 34.1 | 11 | 4 | 43.3 | 227 |
| ETA | 20 | 101.<br>4 | 1288.5 | 19.4 | 52.6 | 10.6 | 2.8 | 36.8 | 170.2 |
| WM | 10 | 120.<br>2 | 824.8 | 30.2 | 73.5 | 9.6 | 4 | 33 | 69.6 |
| SM | 22 | 79.5 | 967.9 | 20.9 | 120.4 | 9.8 | 1.6 | 32.3 | 74.6 |
| CM | 16 | 66.5 | 1206.1 | 14.8 | 118 | 10.4 | 1.6 | 23.8 | 251.7 |
| NM | 8 | 95.3 | 411.4 | 26.5 | 55.4 | 8.8 | 0.8 | 24.5 | 129.1 |

For inflorescence length, we found that Malagasy populations were typically longer, except for CM that was similar to mainland African populations (Table 2). However, the inflorescence length was highly variable and there were few statistically significant differences aside from WM and NM being longer than CM, and WM being longer than all mainland African populations (Table 2; Fig. 3d. There were no significant differences in spikelet or awn length within Madagascar. There was strong correlation among quantitative characters though, with taller plants having longer inflorescences (*r* = 0.59, *p* < 0.001), longer spikelets (*r* = 0.21, *p* = 0.02), and longer awns (*r* = 0.33, *p* < 0.001). There was no clear separation among populations based on traits; although, the first principal component (PC1) does show some separation of WM and NM versus CM (Fig. 3c).

### Distribution Modelling

#### Model training and evaluation

We retrieved a 3,704 occurrence records of *L. simplex* and its synonyms across Africa. This included 2,557 occurrences from GBIF, 1,094 from BIEN and 53 from the field. After removing likely errors and duplicates, we retained 125 occurrence records for Madagascar, 209 for SA, 233 for ETA, and 1354 for all of Africa. Each regional SDM included an equal ratio of pseudo-absences to presence occurrences. *Loudetia simplex* distribution models performed well for SA (AUC = 0.87, TSS = 0.74) and all of Africa (AUC = 0.86, TSS = 0.61), while Madagascar (AUC = 0.68, TSS = 0.40) and ETA (AUC = 0.67, TSS = 0.47) performed moderately.(Supplementary Fig. S1).

Suitable habitats indicated by our models were largely congruent with presence records (Supplementary Fig. S2), but some regions not well-represented in our data were indicated as suitable habitats, such as Sudan and Chad (Fig. 4a). There was qualitative overlap between our African model and the SA model (Fig. 4b) and ETA (Fig. 4c); however, the suitability of some regions were higher in the ETA model, especially in proximity to lake systems in Malawi, Burundi, Rwanda, and Tanzania. The African model indicated a much wider range of *L. simplex* in Madagascar than the Madagascar model (Fig. 4d). The Madagascar model indicated some suitable habitat in regions with no occurrence records and that are not anticipated as the *L simplex* range, such as the high-altitude areas of the northern Sofia region.

#### Variable selection and variable importance

Of the 11 selected variables (Supplementary Table S2), mean annual precipitation was ranked first or second across all four models, with mean annual temperature also ranked highly except for SA (Supplementary Fig. S3). Precipitation seasonality and pH were ranked above temperature for the SA model (Supplementary Fig. S3). Based on the marginal response curves, the Africa model indicated *L. simplex*’s optimal annual precipitation range falls between 1,000 mm and 1,700 mm of annual precipitation with high seasonality and a preference for cooler mean annual temperatures (Fig. 5). These findings are consistent with the Madagascar model based on responsed curves alone, where declines in *L. simplex* presence under increasing precipitation and temperature is more pronounced. Similar patterns were observed for the ETA model, but the trends were not observed in the SA model, where there is typically less precipitation, cooler temperatures, and a different seasonality exposure (Fig. 5).

**Figure 5.**
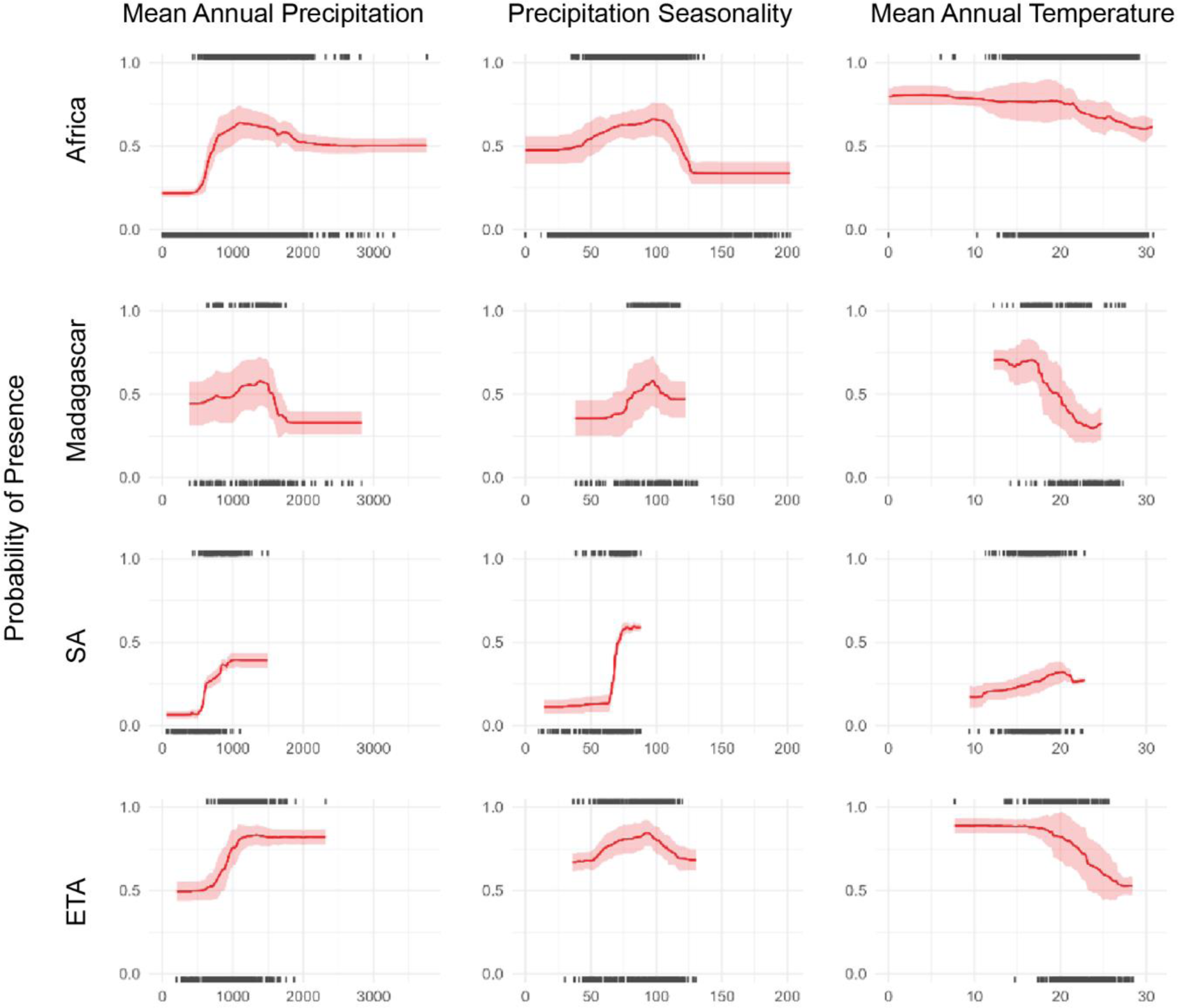
Response curve of the three most important environmental variables across models. From top to bottom, the models are: Africa, Madagascar, SA and ETA. The solid lines are the optimized estimates from the observed data and polygons represent the 95% confidence intervals determined by the permutations. Black tick marks above are presences and tick marks along the bottom of plots are pseudo-absences.

Considering the response curves across all models, *L. simplex* is found across a wide range of precipitation and temperature, and this becomes evident when observing the underlying data. For example, a comparison of the presence records and background pseudoabsences across Africa show there are lower limits to mean annual precipitation for *L*. *simplex* distributions near 500mm, and some occurances approaching 2,000mm (Fig.6a). This corresponds to the underlying regional variation, with drying conditions in SA and wetter conditions in ETA (Fig. 6b). Madagascar has a bimodal distribution that roughly corresponds to precipitation conditions in SA and ETA; although, experiencing more precipitation than the ETA distribution (Fig. 6b). Regarding temperature, the entire distribution is bimodal, reflecting the underlying regional variation. While there are *L. simplex* distributions that stand out from the background pseudo-absences under cooler conditions, *L. simplex* can also be found under warm conditions that approach the limit of the background distribution (Fig. 6c). This reflects the underlying differences of cooler conditions in SA and warmer conditions in ETA (Fig. 6d). Again, Madagascar has a bimodal distribution reflecting both SA and ETA temperature distributions (Fig. 6d)

**Figure 6.**
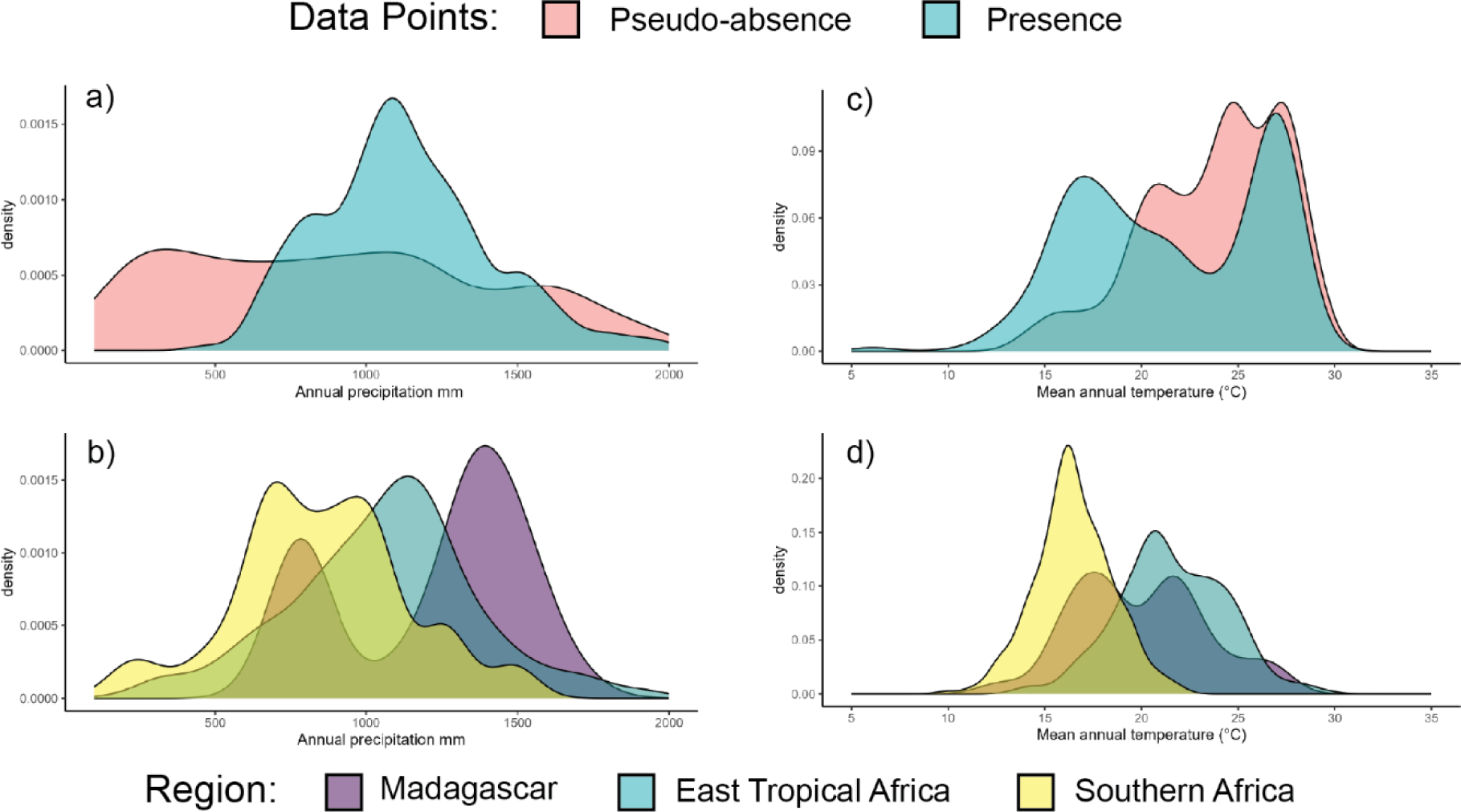
Distributions of mean annual precipitation and temperature. a) Distribution of precipitation among pseudo-absence and presence points across the entire *L. simplex* range. b) Precipitation distributions from presence points used in the Madagascar, East Tropical Africa (ETA), and Southern Africa (SA) models. c) Distribution of temperature among pseudo-absence and presence points across the entire *L. simplex* range. d) Temperature distributions from presence points used in the Madagascar, ETA, and SA models.

#### Niche overlap between Madagascar and mainland Africa

We projected the niches optimized under the SA (Fig. 7a) and ETA (Fig. 7b) models on to Madagascar to quantify niche overlap. The similarity statistics for ranges when masking humid forests was D = 0.33 or I = 0.5 for SA projections (Fig. 7a) and D = 0.45, or I = 0.56 for ETA projections (Fig. 7b). When not masking humid forests, the overlap was notably lower for the SA (D = 0.19 or I = 0.39) and ETA (D = 0.26 or I = 0.44) projections (Supplementary Fig. S4). Despite differences among models, both the SA and ETA projections agree that the Central Highland Plateau is suitable habitat for *L. simplex*. The Madagascar model includes high altitude regions of southwestern and southcentral Madagascar, such as the Ihorombe Plateau and Isalo National Park, that were not captured by the SA or ETA models while both projections suggest almost the entire eastern escarpment is suitable habitat much further north than presently observed in Madagascar (Supplementary Fig. S2).

**Figure 7.**
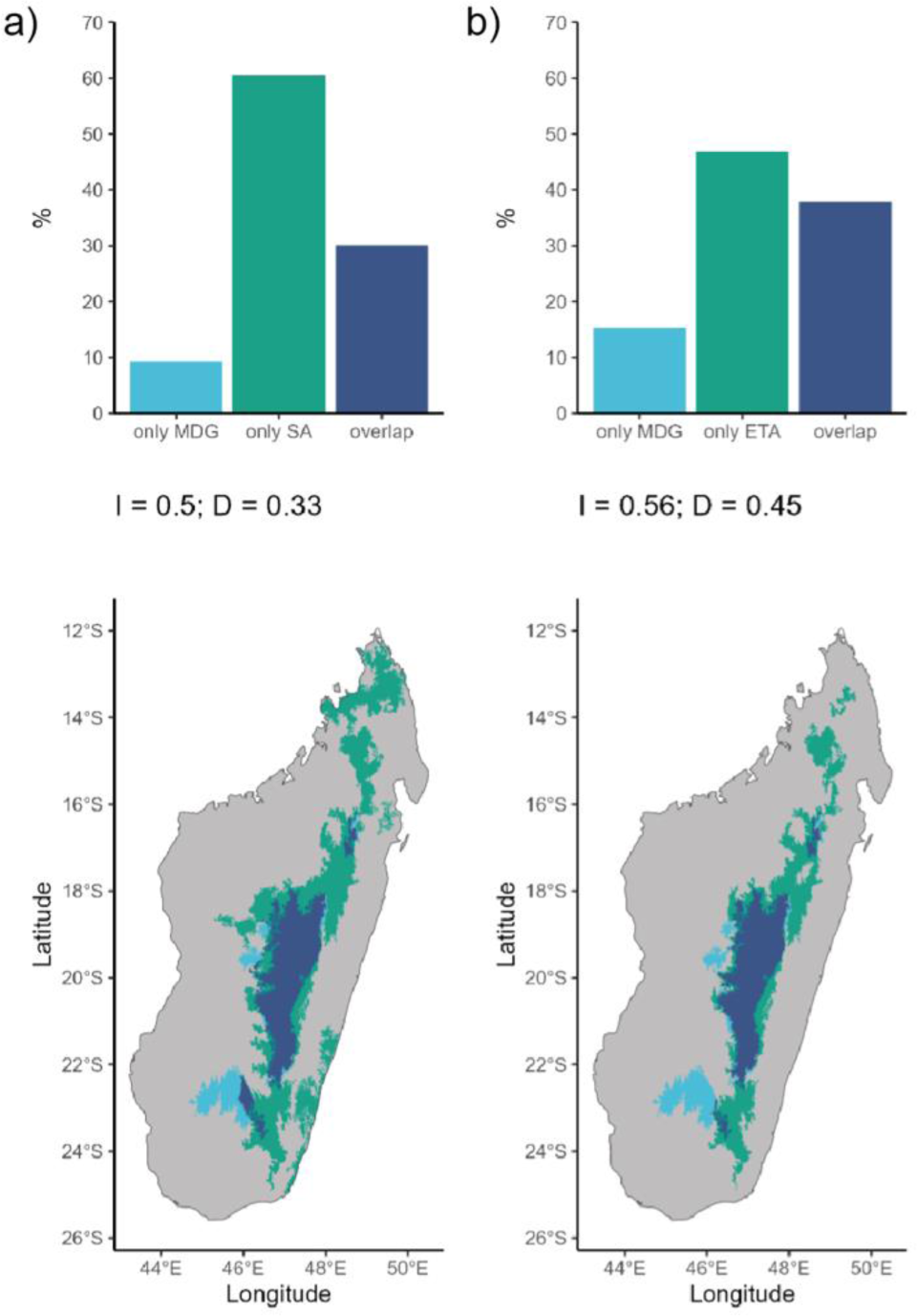
Projection of environmental niches onto Madagascar masking humid forest. The percent of Madagascar predicted as suitable habitat for *Loudetia simplex* and niche overlap statistics for a) the South Africa model and b) the ETA model.

## DISCUSSION

### A single pan-African species

*Loudetia simplex* across mainland Africa and Madagascar likely represents a single species. Although previous hypotheses for a separate Malagasy species or multiple subspecies within Madagascar were made (Bosser 1969), our analyses showed variation within Madagascar was only part of a continuum observed across *L. simplex* more broadly (Fig. 2). Although some morphological variation within Madagascar was observed (Fig. 3c; Table 2), there was no geographic region that could be discriminated from the others and mainland African variation (Fig. 3d). Bosser (1966) recognized *L. simplex* subsp. *stipoides* and *L. madagascariensis* within Madagascar; however, our results suggested the two should be synonymized under *L. simplex* in agreement with Clayton (1974).

The observed morphological variation may be associated with environmental factors as well as genetics. For example, the individuals from western Madagascar and northern Madagascar along with ETA (Fig. 3d) were taller with longer, compacted, inflorescences and short spikelets. The central Madagascar and South African individuals were shorter vegetatively, but with loose inflorescences and longer spikelets (Fig. 3d; Table 2). The variation within Madagascar likely corresponds to the bimodal distributions in precipitation (Fig. 6b) and temperature (Fig. 6d) that reflect the environmental differences between SA and ETA distributions. Therefore, we suggest environment, at least in part, underlies the regional morphological variation within Madagascar.

There is likely a genetic component underlying morphological variation too. A recent microsatellite study (Hagl et al., 2020) showed genetic structure between *L. simplex* from the north of the distribution, overlapping with our NM sample, and the southern extent of the distribution, corresponding to our CM and SM samples (Table 2; Supplementary Table S1). Additionally, Hagl et al., (2020) implied ploidy variation within Madagascar and mainland Africa. Polyploidy is well known to underpin adaptive traits in response to stress (e.g. Fox et al., 2020), such as non-optimal environmental conditions, and plant height in panicoid grasses such as *Zea mays* (Riddle et al., 2006). The precise role of ploidy and genetic variation in the morphological variation of *L. simplex* is not clear at this time, but associations would not be surprising, and these are likely not independent from environmental variation.

### Environmental predictors of Loudetia simplex distributions

At a first glance, *L. simplex* appears to prefer mean annual temperatures between 10C and 20C, and mean annual precipitation between 1,000 mm and 1,700 mm with high seasonality (i.e. a wet and dry season; Fig. 5). These trends are typical features of grassland ecosystems, especially the quantity and seasonality of precipitation that allows grasslands to produce biomass and then desiccate, driving fires in the dry seasons (Lehmann et al., 2011; Alvarado et al., 2020). However, high regional variation underlies precipitation and temperature (Fig. 6). *Loudetia simplex* can be found under higher precipitation conditions up to 2,000 mm in both Madagascar and ETA (Fig. 6b), but this reflects the limitations of the background environment (Fig. 6a). A previous study suggested some Malagasy grasslands were anthropogenic because they received more precipitation than expected from other mainland African grasslands (Joseph et al., 2022b). However, as demonstrated here, focusing only on African-wide marginal response curves can place unreasonable restrictions on discourse about which grasslands are anthropogenic and which are not, when the underlying environmental conditions for a widespread species such as *L. simplex* varies greatly with region (Fig. 6)

The environmental models nevertheless struggle to predict where *L. simplex* occurs in Madagascar. For example, the SA and ETA models without masking suggested suitable habitats in Madagascar’s humid forests (Supplementary Fig. S4), but this fails to account for the demographic dynamics and physiological constraints, such as shading, between closed-canopy and open ecosystems (Staver 2018; Aleman & Staver 2018, Charles-Dominique et al., 2018). When masking the humid forests, there is far more congruence between the projected distributions, but at best about 50% overlap (depending on the statistic) between the SA and ETA models versus the Madagascar niche (Fig. 7). A notable feature omitted from our models was fire. The scale and frequency of fire is important in determining where grasslands occur (Phelps et al., 2022), and this should be especially true of the distribution of *L. simplex* that has adaptive traits for fire-driven ecosystems (Solofondranohatra et al., 2020). Thus, there may be avenues for improving future distribution models, but even limiting our inferences to environmental variables strongly supports Madagascar’s Central Highlands as suitable habitat for *L. simplex* where it can be found in high abundance.

### Implications for Madagascar’s grasslands

Resolving taxonomic uncertainties regarding *L. simplex* in Madagascar has helped us place Malagasy grasslands in context with similar mainland African ecosystems. Much of the Central Highlands were recovered as suitable habitat, which extends from Anjozorobe, in the vicinity of the Ankafobe and Ambohitantely protected areas represented by NM in the morphological analyses (Fig. 2; Supplementary Table S1), to the Itremo protected area and Isalo Massif represented by SM morphological sample. This identified area likely represents pre-human *L. simplex* dominated grasslands with natural fire cycles. There is strong support for pre-human grasslands in Madagascar (Bond et al., 2008; Vorontsova et al., 2016; Crowley et al., 2021; Bond et al., 2023), but it is also well-accepted that these open biomes have likely expanded through the Pleistocene and in response to anthropogenic activities (Burney 1987; Gasse & van Campo 1998). Determining which grasslands represent recent expansions is difficult, especially since our distribution models are largely driven by climatic variables, and there are certainly interactions between climate, soil, fire, herbivory, and vegetation that shape the *L. simplex* distribution (Lehmann et al., 2014). However, connecting morphology, distribution models, and genetics (e.g. Hagl et al., 2020) of *L. simplex* may be helpful for identifying recently expanded grasslands. These efforts will be facilitated by our recommendation to synonymize *L. simplex* subsp. *stipoides* and *L. madagascariensis* under *L. simplex*.

## Supporting information

Measurement data

## ACKNOWLEDGEMENTS

TOMA was supported by the Royal Botanic Garden Edinburgh and an Emily Holmes Memorial Scholarship courtesy of the Amar-Franses & Foster-Jenkins Trust. This project has received funding from the European Union’s Horizon 2020 research and innovation programme under the Marie Sklodowska-Curie grant agreement No. 101026923, awarded to GPT and MSV. New field collections were conducted with permits 049/22/MEDD/SG/DGGE/DAPRNE/SCBE.Re and 050/22/MEDD/SG/DGGE/DAPRNE/SCBE.Re issued to GPT by the Madagascar Ministry of Environment, Director of Protected Areas, Natural Renewable Resources, and Ecosystems (MEDD DAPRNE). The authors are grateful to the MEDD DAPRNE staff that made collections possible. Collections would not have been possible without the administrative assistance Landy R. Ralaiveloarisoa, Bodoarisoa Mbolatahiana, and colleagues at the Kew Madagascar Conservation Centre (KMCC). Fieldwork was successful only with the guidance of Roger Rajaonarison from KMCC. Herbarium access was granted by the curators of the Parc Botanique et Zoologique de Tsimbazaza herbarium, Madagascar, and the Royal Botanic Gardens, Kew herbarium, UK.

## AUTHOR CONTRIBUTION

TOMA, CERL, MR, GPT, and MSV conceived the study. TOMA, FR, JR, and GPT conducted fieldwork. FR and HR secured permits. TOMA made morphological measurements. TOMA and JW assembled occurrence data. TOMA, GPT, and JW performed analyses. TOMA, GPT, and JW wrote the first draft. All authors revised and approved the final manuscript.

## DATA AVAILABILITY STATEMENT

Morphological data is available in Supplementary Table S1. Scripts for recreating SDM analyses, including the collection of occurrence records from online databases, are available on TOMA’s GitHub page (https://github.com/AlmaryT/Loudetia_SDM).

## REFERENCES

Abatzoglou, J.T., Dobrowski, S.Z., Parks, S.A., & Hegewisch, K.C. (2018). TerraClimate, a high-resolution global dataset of monthly climate and climatic water balance from 1958-2015. Scientific Data, 5, 170191. 10.1038/sdata.2017.191

Aleman, J.V., & Staver, A.C. (2018). Spatial patterns in the global distributions of savanna and forest. Global Ecology and Biogeography, 27(7), 792–803. 10.1111/geb.12739

Ali, J.R., & Aitchison, J.C. (2008). Gondwana to Asia: Plate tectonics, paleogeography and the biological connectivity of the Indian sub-continent from the Middle Jurassic through latest Eocene (166-35 Ma). Earth-Science Reviews, 88, 145–166.

Alvarado, S.T., Andela, N., Silva, T.S.F., & Archibald, S. (2020). Thresholds of fire response to moisture and fuel load differ between tropical savannas and grasslands across continents. Global Ecology and Biogeography, 29(2), 331–433. 10.1111/geb.13034

Amatulli G., McInerney D., Sethi T., Strobl, P., & Domisch, S. (2020). Geomorpho90m, empirical evaluation and accuracy assessment of global high-resolution geomorphometric layers. Scientific Data, 7, 162. 10.1038/s41597-020-0479-6

Antonelli, A., Smith, R. J., Perrigo, A. L., Crottini, A., Hackel, J., Testo, W., Farooq, H., Torres Jiménez, M. F., Andela, N., Andermann, T., Andriamanohera, A. M., Andriambololonera, S., Bachman, S. P., Bacon, C. D., Baker, W. J., Belluardo, F., Birkinshaw, C., Borrell, J. S., Cable, S., … Ralimanana, H. (2022). Madagascar’s extraordinary biodiversity: Evolution, distribution, and use. Science, 378(6623). 10.1126/science.abf0869

Bond, W. J., Silander, J. A., Ranaivonasy, J., & Ratsirarson, J. (2008). The antiquity of Madagascar’s grasslands and the rise of C4 grassy biomes. Journal of Biogeography, 35(10), 1743–1758. 10.1111/j.1365-2699.2008.01923.x

Bond, W. J., Silander, J. A., & Ratsirarson, J. (2023). Madagascar’s grassy biomes are ancient and there is much to learn about their ecology and evolution. Journal of Biogeography, 50(3), 614–621. 10.1111/jbi.14494

Bosser, & Jean. (1966). Notes sur les graminées de Madagascar : 5-Le genre Loudetia Hochst. ex. Steud.

Bosser, J. (1969). GRAMINES DES PATURAGES ET DES CULTURES A MADAGASCAR.

Brummitt, R. K. (2001). World geographic scheme for recording plant distributions, edition 2. Carnegie Mellon University (Pittsburgh): Hunt Institute for Botanical Documentation. http://rs.tdwg.org/wgsrpd/doc/data/

Buerki S., Devey, D.S., Callmander, M.W., Phillipson, P.B., Forest, F. 2013. Spatio-temporal history of the endemic genera of Madagascar. Botanical Journal of the Linneaen Society, 171(2), 304–329. 10.1111/boj.12008

Burney, D. A. (1987). Late Holocene vegetational change in central Madagascar. Quaternary Research, 28(1), 130–143. 10.1016/0033-5894(87)90038-X

Burney, D.A., Burney, L.P., Godfrey, L.R., Jungers, W.L., Goodman, S.M., Wright, H.T. and Jull, A.T., 2004. A chronology for late prehistoric Madagascar. Journal of Human Evolution, 47(1-2), pp.25–63.

Callmander, M.W., Phillipson, P.B., Schatz, G.E., Andriambololonera, S., Rabarimanarivo, M., Rakotonirina, N., Raharimampionona, J., Chatelain, C., Gautier, L., & Lowry, P.P. (2011). The endemic and non-endmemic vascular flora of Madagascar updated. Plant Ecology and Evolution, 144, 121–125.

Charles-Dominique, T., Midgley, G. F., Tomlinson, K. W., & Bond, W. J. (2018). Steal the light: shade vs fire adapted vegetation in forest–savanna mosaics. New Phytologist, 218(4), 1419–1429.

Clayton WD. 1974. Loudetia. In: Clayton WD, Phillips SM, Renvoize SA, eds. Flora of Tropical East Africa. Gramineae 2. London: Crown Agents for Oversea Governments and Administrations, pp. 415–421.

Crottini, A., Madsen, O., Poux, C., Strauß, A., Vietes, D.R., & Vences, M. (2012). Vertebrate time-tree elucidates the biogeographic pattern of a major biotic change around the K-T boundary in Madagascar. Proceedings of the National Academy of Sciences USA, 109, 5358–5363.

Crowley, B. E., Godfrey, L. R., Hansford, J. P., & Samonds, K. E. (2021). Seeing the forest for the trees—and the grasses: revisiting the evidence for grazer-maintained grasslands in Madagascar’s Central Highlands. Proceedings of the Royal Society B, 288(1950), 20201785. 10.1098/rspb.2020.1785

Edwards, E.J., & Smith, S.A. (2010). Phylogenetic analyses reveal the shady history of C_4_ grasses. Proceedings of the National Academy of Sciences USA, 107, 2532–2537.

Fox, D.T., Soltis, D.E., Soltis, P.S., Ashman, T.-L., Van de Peer, Y. (2020). Polyploidy: A biological force from cells to ecosystems. Trends in Cell Biology, 30(9), 688–694. 10.1016/j.tcb.2020.06.006

Gasse, F., & Van Campo, E. (1998). A 40,000-yr pollen and diatom record from Lake Tritrivakely, Madagascar, in the southern tropics. Quaternary Research, 49(3), 299–311. 10.1006/qres.1998.1967

GBIF.org (28 November 2022) GBIF Occurrence Download 10.15468/dl.p4yjv2

Goodman, S. M., Raherilalao, M. J., & Wohlauser, S. (2018). Les aires protégées terrestres de Madagascar : leur histoire, description et biote

Gorelick, N., Hancher, M., Dixon, M., Ilyushchenko, S., Thau, D., & Moore, R. (2017). Google Earth Engine: Planetary-scale geospatial analysis for everyone. Remote Sensing of Environment, 202, 18–27. 10.1016/j.rse.2017.06.031

Hackel, J., Vorontsova, M. S., Nanjarisoa, O. P., Hall, R. C., Razanatsoa, J., Malakasi, P., & Besnard, G. (2018). Grass diversification in Madagascar: In situ radiation of two large C3 shade clades and support for a Miocene to Pliocene origin of C4 grassy biomes. Journal of Biogeography, 45(4), 750–761. 10.1111/jbi.13147

Hagl, P. A., Gargiulo, R., Fay, M. F., Solofondranohatra, C., Salmona, J., Suescun, U., Rakotomalala, N., Lehmann, C. E. R., Besnard, G., Papadopulos, A. S. T., & Vorontsova, M. S. (2020). Geographical structure of genetic diversity in Loudetia simplex (Poaceae) in Madagascar and South Africa. Botanical Journal of the Linnean Society, 1–19. 10.1093/botlinnean/boaa098/6050729

Hijmans, R.J., Cameron, S.E., Parra, J.L., Jones, P.G., & Jarvis, A. (2005). Very high resolution interpolated climate surfaces for global land areas. International Journal of Climatology, 25(15), 1965–1978. 10.1002/joc.1276

Hijmans, R.J., Phillips, S., Leathwick, J., Elith, J. (2022). dismo: species distribution modeling. R package version 1.3-9. Retrieved from https://CRAN.R-project.org/package=dismo

Jarvis, A., Reuter, H.I., Nelson, A., & Guevara E. (2008). Hole-filled SRTM for the globe Version 4, available from the CGIAR-CSI SRTM 90m Database: https://srtm.csi.cgiar.org.

Joseph, G.S., Rakotoarivelo, A.R., & Seymour, C.L. (2022a). Tipping points induced by palaeo-human impacts can explain presence of savannah in Malagasy and global systems where forest is expected. Proc Biol Sci, 289,20212771.

Joseph, G.S., Rakotoarivelo, A.R., & Seymour, C.L. (2022b). How expansive were Malagasy Central Highland forests, ericoids, woodlands and grasslands? A multidisciplinary approach to a conservation conundrum. Biological Conservation, 261(1), 109282. 10.1016/j.biocon.2021.109282

Lehmann, C.E.R., Anderson, T.M., Sankaran, M., Higgins, S.I., Archibald, S., Hoffmann, W.A., Hanan, N.P., Williams, R.J., Fensham, R.J., Felfili, J., Hutley, L.B., Ratnam, J., San Jose, J., Montes, R., Franklin, D., Russell-Smith, J., Ryan, C.M., Durigan, G., Hiernaux, P., Haidar, R., Bowman, D.M.J.S., & Bond, W.J. (2014). Savanna vegetation-fire-climate relationships differ among continents. Science, 343(6170), 548–552. 10.1126/science.1247355

Lehmann, C.R., Archibald, S.A., Hoffmann, W.A., Bond, W.J. (2011). Deciphering the distribution of the savanna biome. New Phytologist, 191(1), 197–209. 10.1111/j.1469-8137.2011.03689.x

Leroy, B., Meynard, C.N., Bellard, C., & Courchamp, F. (2015). Virtualspecies, an R package to generate virtual species distributions. Ecography, 39(6), 599–607. 10.1111/ecog.01388

Miller, M.A, Shepherd, K.D., Kisitu, B., & Collinson, J. (2021). iSDAsoil: The first continent-scale soil property map at 30 m resolution provides a soil information revolution for Africa. PLoS Biology, 19(11), e3001441. 10.1371/journal.pbio.3001441

Moat, J., & Smith, P. 2007. Atlas of the Vegetation of Madagascar (Atlas de la Végétation de Madagascar). Royal Botanic Gardens, Kew.

Myers, N., Mittermeier, R.A., Mittermeier, C.G., da Fonseca, G.A.B, & Kent J. 2000. Biodiversity hotspots for conservation priorities. Nature, 403, 853–858.

Phelps, L.N., Andela, N., Gravey, M., Davis, D.S., Kull, C.A., Douglass, K., & Lehmann, C.E.R. (2022). Madagascar’s fire regimes challenge global assumptions about landscape degradation. Global Change Biology, 28(23), 6944–6960. 10.1111/gcb.16206

Ralimanana, H., Perrigo, A. L., Smith, R. J., Borrell, J. S., Faurby, S., Rajaonah, M. T., Randriamboavonjy, T., Vorontsova, M. S., Cooke, R. S. C., Phelps, L. N., Sayol, F., Andela, N., Andermann, T., Andriamanohera, A. M., Andriambololonera, S., Bachman, S. P., Bacon, C. D., Baker, W. J., Belluardo, F., … Antonelli, A. (2022). Madagascar’s extraordinary biodiversity: Threats and opportunities. Science, 378(6623). 10.1126/science.adf1466

Riddle, N.C., Kato, A., & Birchler, J.A. (2006). Genetic variation or the response to ploidy change in *Zea mays* L. Theoretical and Applied Genetics, 114(1), 101–111. 10.1007/s00122-006-0414-z

Schoener, T.W. (1968). The *Anolis* lizards of Bimini: Resource partitioning in a complex fauna. Ecology, 49(4), 704–746. 10.2307/1935534

Šlenker, M., Koutecký P., & Marhold K. 2022. MorphoTools2: an R package for multivariate morphometric analysis. Bioninformatics, 38(10), 2954–2955. 10.1093/bioinformatics/btac173

Solofondranohatra, C. L., Vorontsova, M. S., Hempson, G. P., Hackel, J., Cable, S., Vololoniaina, J., & Lehmann, C. E. R. (2020). Fire and grazing determined grasslands of central Madagascar represent ancient assemblages: Grasslands are shaped by disturbance. Proceedings of the Royal Society B: Biological Sciences, 287(1927). 10.1098/rspb.2020.0598

Staver, A.C. (2018). Prediction and scale in savanna ecosystems. New Phytologist, 219(1), 52–57. 10.1111/nph.14829

Velazco, S.J.E., Rose, M.B., de Andrade, A.F.A., Minoli, I., Franklin, J. (2022). FLEXSDM: An R package for supporting a comprehensive and flexible species distribution modelling workflow. Methods in Ecology and Evolution, 13(8), 1661–1669. 10.1111/2041-210X.13874

Vorontsova, M. S., Besnard, G., Forest, F., Malakasi, P., Moat, J., Clayton, W. D., Ficinski, P., Savva, G. M., Nanjarisoa, O. P., Razanatsoa, J., Randriatsara, F. O., Kimeu, J. M., Quentin Luke, W. R., Kayombo, C., & Peter Linder, H. (2016). Madagascar’s grasses and grasslands: Anthropogenic or natural? Proceedings of the Royal Society B: Biological Sciences, 283(1823). 10.1098/rspb.2015.2262

Warren, D.L., Glor, R.E., Turelli, M. (2008). Environmental niche equivalency versus conservatism: quantitative approaches to niche evolution. Evolution, 62(11),2868–2883. 10.1111/j.1558-5646.2008.00482.x

Zizka, A., Silvestro, D., Andermann, T., Azevedo, J., Ritter, C.D., Edler, D., Farooq, H., Herdean, A., Ariza, M., Scharn, R., Svantesson, S., Wengström, N., Zizka, V., & Antonelli, A. (2019). CoordinateCleaner: Standardized cleaning of occurrence records from biological collection databases. Methods in Ecology and Evolution, 10(5), 744–751. 10.1111/2041-210X.13152

