## Supplementary material for "The grass that built the Central Highland of Madagascar: environmental niches and morphological diversity of *Loudetia simplex*"

### Morphological and ecological diversity of *Loudetia simplex* in Madagascar and in Africa

SUPPLEMENTARY FIGURES

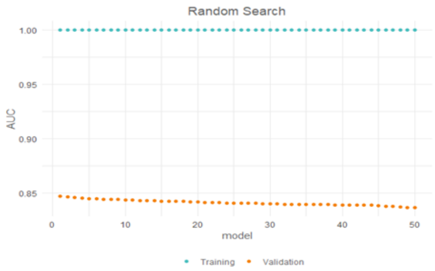


*Africa*

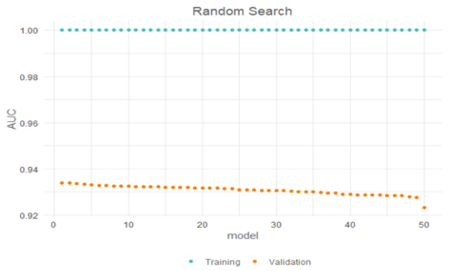


*South Africa*

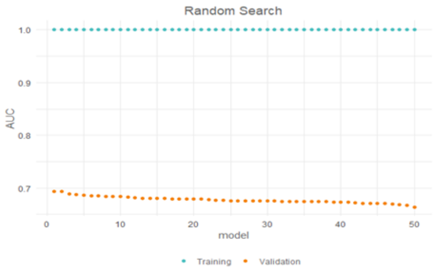


*East tropical Africa*

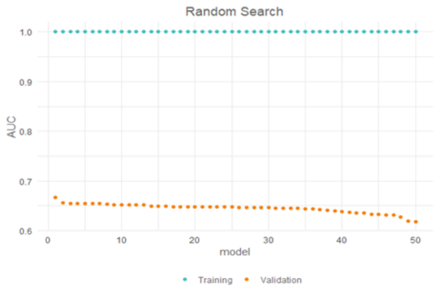


*Madagascar*

Variable importance for each locality

| | AUC | TSS |
|---|---|---|
| AFR | 0.84 | 0.57 |
| ETA | 0.68 | 0.37 |
| SA | 0.92 | 0.82 |
| MDG | 0.64 | 0.38 |

**Fig. S1 — AUC and TSS values for distribution models by region.**

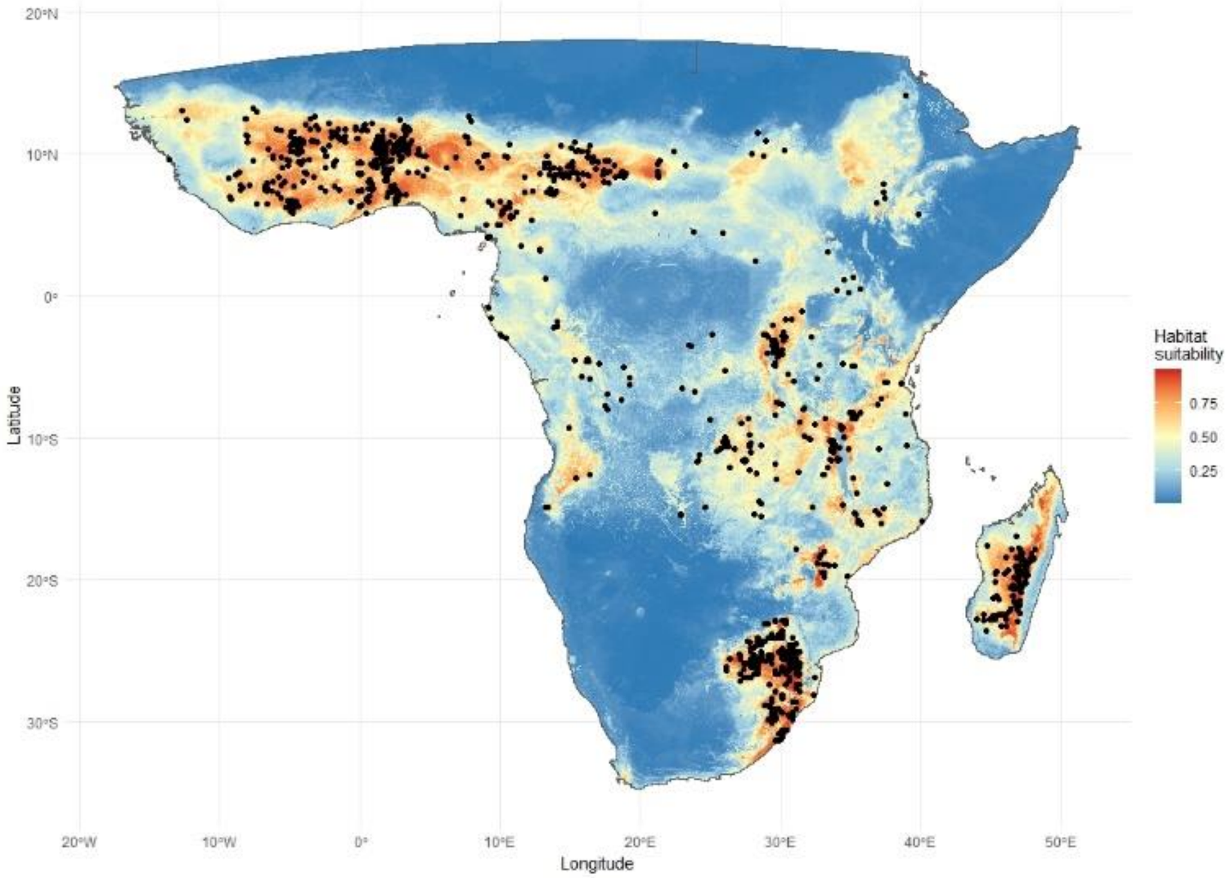


**Fig. S2 — *L. simplex* occurrence and habitat suitability.** Individual points represent occurrence records retained after filtering. Habitat suitability shown is based on the Africa model.

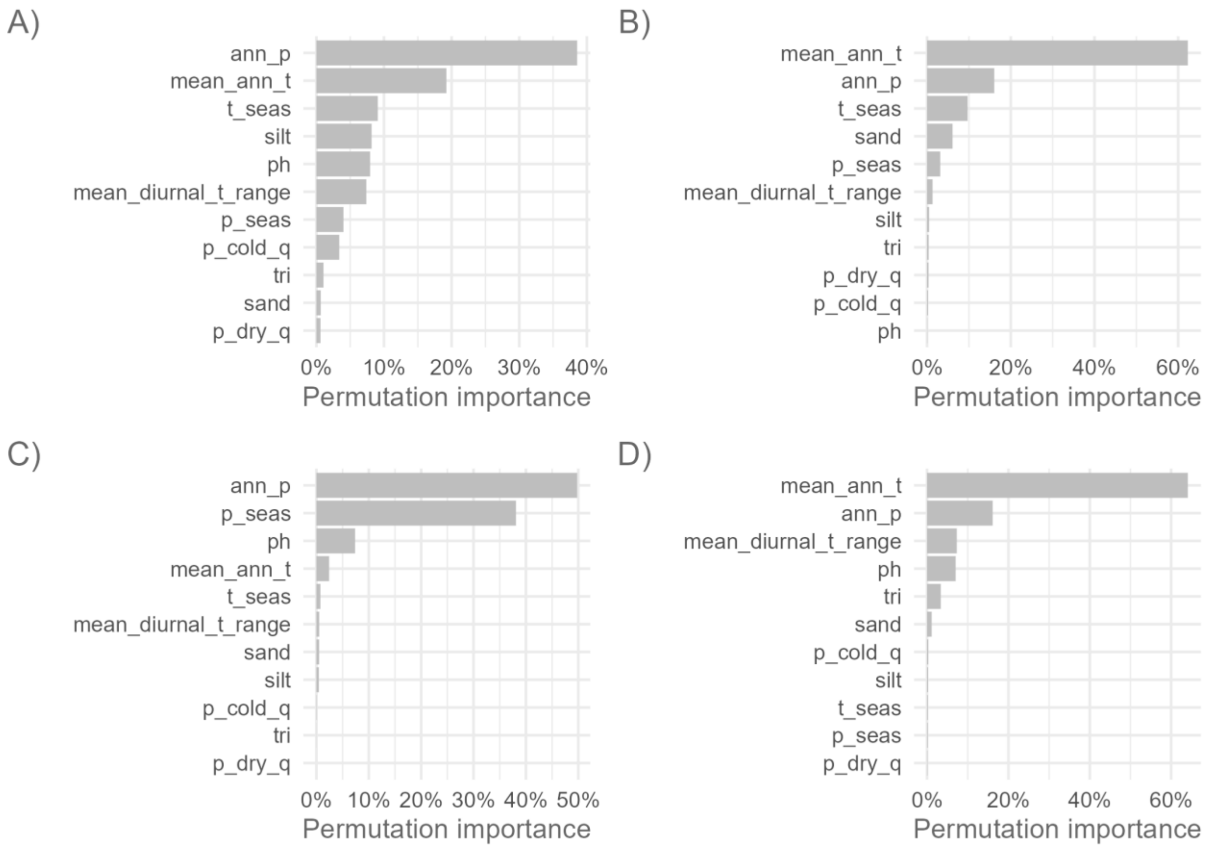


**Fig. S3 — Ranked variable importance.** Percent of model weight explained by environmental parameters for A) Africa, B) Madagascar, C) South Africa, and D) east tropical Africa (ETA).

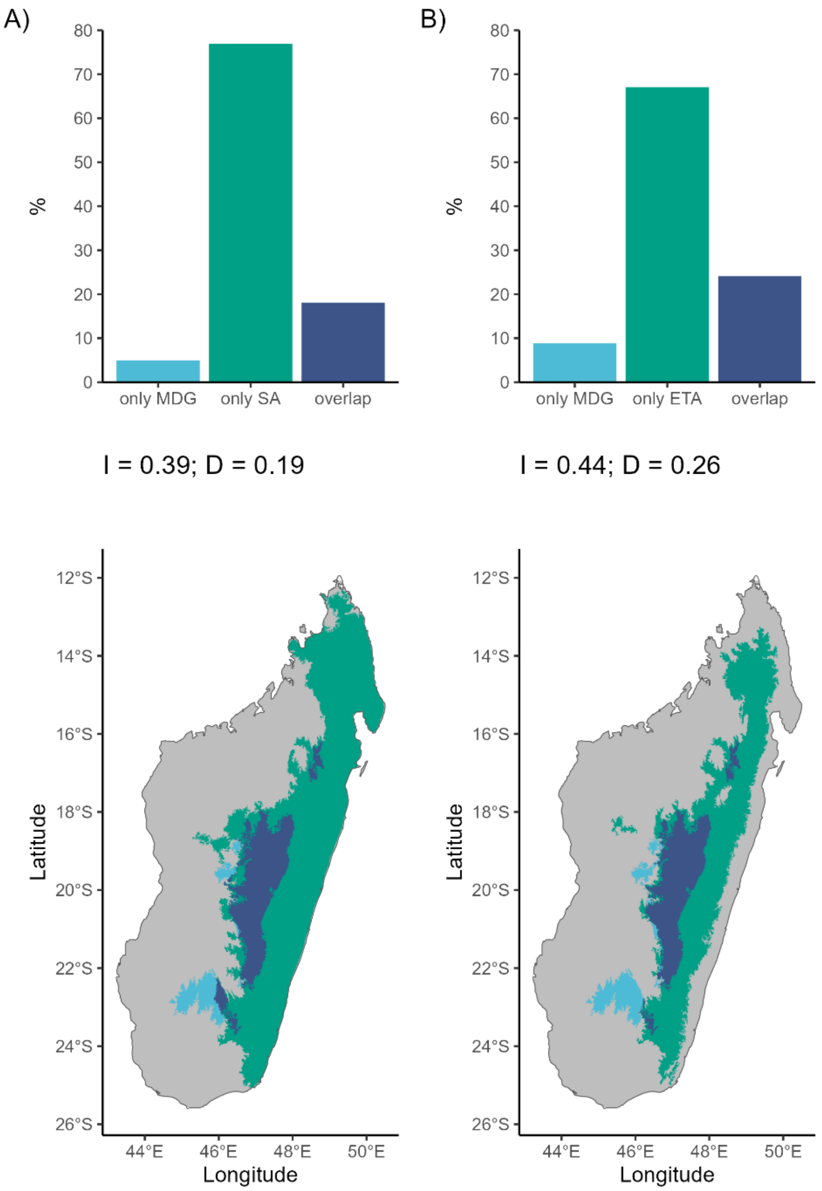


**Fig. S4 — Projection of environmental niches onto Madagascar without masking humid forests**. The percent of Madagascar predicted as suitable habitat for *Loudetia simplex* and niche overlap statistics for A) the South Africa model and B) the ETA model.
